# Fungal succession on the decomposition of three plant species from a Brazilian mangrove

**DOI:** 10.1101/2021.06.09.447690

**Authors:** Marta A. Moitinho, Josiane B. Chiaramonte, Laura Bononi, Thiago Gumiere, Itamar S. Melo, Rodrigo G. Taketani

**Affiliations:** Laboratory of Environmental Microbiology, Brazilian Agricultural. Research Corporation, EMBRAPA Environment, SP 340. Highway—Km 127.5, Jaguariúna, SP 13820-000, Brazil; College of Agriculture Luiz de Queiroz, University of São Paulo, Pádua Dias Avenue, 11, Piracicaba, SP 13418-900, Brazil; Institut National de la Recherche Scientifique, Centre Eau Terre Environnement. 490, rue de la Couronne, Quebec City, QC, G1K 9A9, Canada; CETEM, Centre for Mineral Technology, MCTIC Ministry of Science, Technology, Innovation and Communication, Av. Pedro Calmon, 900, Cidade Universitária, Ilha do Fundão, Rio de Janeiro, Brazil, ZC 21941-908

**Author notes:** Correspondance author: College of Agriculture Luiz de Queiroz, University of São Paulo, Pádua Dias Avenue, 11, Piracicaba, SP 13418-900, Brazil.

**Keywords:** fungal dynamics, Mangrove plant degradation, succession, Microcosms

## Abstract

Leaf decomposition is the primary process in release nutrients in the dynamic mangrove habitat, supporting the ecosystem food webs. On most environments, fungi are an essential part of this process. However, due to the peculiarities of mangrove forests, this group is currently neglected. Thus, this study tests the hypothesis that fungal community display an specific succession pattern in different mangrove species. A molecular approach was employed to investigate the dynamics of the fungal community during the decomposition of three common plant species (*Rhizophora mangle*, *Laguncularia racemosa*, and *Avicennia schaueriana*) from the mangrove habitat located at the southeast of Brazil. Plant material was the primary driver of fungi communities but time also was marginally significant for the process, and evident changes in the fungal community during the decomposition process were observed. The five most abundant classes common to all the three plant species were *Saccharomycetes*, *Sordariomycetes*, *Tremellomycetes*, *Eurotiomycetes*, and *Dothideomycetes*, all belonging to the Phylum Ascomycota. *Microbotryomycetes* class were shared only by *A. schaueriana* and *L. racemosa*, while *Agaricomycetes* class were shared by *L. racemosa* and *R. mangle*. The class *Glomeromycetes* were shared by *A. schaueriana* and *R. mangle*. The analysis of the core microbiome showed that *Saccharomycetes* was the most abundant class. In the variable community, Sordariomycetes was the most abundant one, mainly in the *Laguncularia racemosa* plant. The results presented in this work shows a specialization of the fungal community regarding plant material during mangrove litter decomposition.

## 1. Introduction

Mangroves are unique ecosystems composed by autochthonous vegetation of marine plants, which grows in marshy tidal areas of estuaries and coastal shorelines of the tropics and subtropics (Holguin et al. 2001; Raghukumar 2017). The mangrove vegetation adapted morphologically and physiologically to the fluctuating conditions of salinity, temperature and, in the sediment, low oxygen concentrations (Sebastianes et al. 2013; Raghukumar 2017). A critical aspect of mangroves lies in its export of organic matter to adjacent coastal areas (Schaeffer-Novelli et al. 1990; Alongi 1994; Holguin et al. 2001, 2006; Baskaran et al. 2012; Taketani et al. 2018). Supporting estuarine food webs and commercial fisheries (Holguin et al. 2001).

The mangrove is an environment where fungi can thrive due to a large amount of litter produced by its vegetation (Raghukumar 2017). Besides, the metabolic versatility of this microorganisms allows them to exploit many different niches which renders them as significant components of these ecosystems, acting as decomposers, mutualists, and pathogens (Schmit and Mueller 2007). However, despite the diversity of approximately 1.5 million species (Hawksworth, D L 2004), we have identified a minority of these species (Schmit and Mueller 2007) and even less in marine habitats (Valderrama et al. 2016).

Fungi play an essential role in the decomposition of lignocellulosic material in mangroves (Sebastianes et al. 2013). During these processes, fungi are considered one of the most active microorganisms, more active than bacteria, mainly in the early stages of the process (Pascoal and Cassio 2004; Marano et al. 2011; Raghukumar 2017). They inhabit dead wood, prop roots, leaves, and viviparous seedlings, and their dynamics in this habitat depends on the physical and chemical properties of the substrate (Raghukumar 2017). Thus, it is of great importance to have additional information about fungi diversity and distributions to improve the predictions about the diversity of fungal communities associated with different plant species around the world (Schmit and Mueller 2007).

Succession is the process of change in community composition over time in an ordinated manner (Marano et al. 2011; Moitinho et al. 2018). Studies of fungi decomposition showed this process many marine ecosystems (Tan et al. 1989; Ananda and Sridhar 2004; Maria et al. 2006). For the fungal communities, the decomposition process of leaf litter generally occurs in three distinct phases: early, intermediate, and late decomposers (Baldrian et al. 2012). However, the degradation of leaves in mangroves is dynamic and generally occurs in a matter of a few weeks (Raghukumar 2017) which can alter this organized process. Little is known about the role of fungi during the decomposition process in mangroves ecosystems using next-generation sequencing. Therefore, the aim of this study was to evaluate and compare the diversity of fungi community during the decomposition process of three mangrove plant species from Cananéia mangrove, Brazil. We set the assays using microcosms conditions to follow up the decomposition process of the three plant species during four different periods. Therefore, this study hypothesized that during leaf decomposition in Cananéia mangrove, different plant species select for a specific community structure along the process and this selection reflect on differences in the role of the observed populations.

## 2. Material and Methods

### 2.1. Site Description and Collected Material

Samples were collected from one mangrove forest in the city of Cananéia in São Paulo state, Brazil. The mangrove located in Cananéia (Can) (25° 05′ 03″ S–47° 57′ 75″ W) a pristine area with little human influence. Samples were taken during the low tide. Most mangroves of the south coast of Brazil have homogeneous vegetation composed of *Rhizophora mangle, Laguncularia racemosa, and Avicennia schaueriana.*

Fresh mature leaves that did not present any sign of lichen or lesion of any sort were carefully collected directly from these three plant species (avoiding contact with sediment, water or human DNA). Five kilograms of the top 5 cm of sediments were collected in each mangrove plant species, and 5 l of brackish estuarine water was also sampled in the proximities of the sampling sites. The material was immediately placed in sterile bags and transported to the laboratory for immediate processing and microcosm implementation.

### 2.2. Microcosm’s Construction

The experiment was conducted in a completely randomized design consisting of 3 plant species X 4 time periods X 3 repetitions totalizing 36 samples. Microcosms were built using 50 ml penicillin flasks (50 mm diameter) to evaluate the fungal dynamic throughout the organic matter decomposition process. Microcosms were constructed in three replicates, and each one contained 20 g of sediment, eight disks of leaves randomly selected (1.7 cm diameter) from one of the three plant species without any previous treatment on order to decrease external influences in the treatments and simulate mangrove ecosystems and 2 ml of brackish water was placed on top. They were incubated at 24 °C and monitored during 60 days with subsamples taken at 7, 15, 30, and 60 days. At each sampled date, four leaf disks were collected and used to extract the DNA (Moitinho et al. 2018). With the intent to have the acclimatization and stabilization in the microcosms, by the mixture of the fresh leaves along the sediment and water simulating the environmental conditions found in the mangrove sediment, no subsample was taken at time zero, and the first sample in day seven was considered as a parameter.

### 2.3. Fungi communities’ analysis using ITS region sequencing

The samples of decaying leaves were submitted to DNA extraction using the PowerSoil DNA Isolation Kit (MoBio Carlsbad, USA) according to the manufacturer’s instructions. These extractions were made by transferring 0.25 g of plant material directly from the microcosms into the PowerBead tubes provided in the Power Soil. Quality and quantity of the extracted DNA were evaluated by Nanodrop spectrophotometer (Thermo Scientific 2000 spectrophotometer) and in 2.4% agarose gel.

The samples were PCR amplified using the primers set ITS1f (Gardes and Bruns 1993) and ITS2 (White et al. 1990), along with tags of five nucleotides with the forward primer to distinguish the samples, and to generate the amplicons of the ITS rRNA gene. The amplification mix contained 18 μl of water; 2.5 μl of PCR buffer; 3.3 μl of MgCl_2_; 0.5 μl od dNTPs; 0.5 μl of Taq DNA Polymerase (Invitrogen, São Paulo, Brazil); 0.25 μl of each primer and finally 0.5 μl of DNA. Cycling conditions were an initial denaturation of 94°C for 3 min; 30 cycles of 94°C 30 s; 57°C for 45 s; and 72°C for 1 min; and a final 2 min extension at 72°C.

The PCR product was purified with the Promega Wizard® SV Gel and PCR Clean-Up System Kit, quantified by Qubit 2.0 Fluorometer (Life Technologies) and Sequencing was performed on an Ion Torrent PGM system of Life Technologies using the Ion 316™ Chip. The enrichment phase was performed by using the One-Touch 2 device with the Ion Sequencing 400 Kit according to the manufacturer’s instruction (Life Technologies) (Moitinho et al. 2018). Raw sequencing data obtained from the PGM were processed using QIIME software (Quantitative Insights Into Microbial Ecology) following the tutorial described previously.

### 2.4. Processing of the sequences

Raw sequencing data obtained from the PGM were processed using QIIME software (Quantitative Insights Into Microbial Ecology) (Caporaso et al. 2010) following a modified version of the 454 Overview Tutorials using the UNITE database (Kõljalg et al. 2013). The identification and removing of chimaeras were made with the following commands “identify_chimeric_seqs.py” and “filter_fasta.py” respectively. The Otus were picked with the command “pick_otus.py”, and the selection of the representative Otus was made with the “pick_rep_set.py” command. The taxonomies were assigned with “assign_taxonomy.py”, and the OTU tables were assembled with “make_otu_table.py”. The normalization values were obtained with the command “biom summarize-table”.

After the initial filtering steps, we obtained 314786 million reads with an average of 8744.056 reads per library (min. 4196.0 and max. 18777.0). After removing unclassified microorganisms not belonging to the Fungi reign, the reads count dropped to 11646 taxa distributed in 36 samples and seven variables (3 plant species X 4 time periods X 3 repetitions). After the rarefaction step, the taxonomic table remained with 6378 taxa among five taxonomic ranks that were used in downstream analyses.

### 2.5. Statistical analyses

We evaluated the composition of the whole community and also the core and variable portions of fungi communities during the decomposition process of three plant species from Cananéia Mangrove. To improve the understanding from the 26739 representative OTUs picked, only 11648 were classified in the kingdom Fungi and were retained for analysis. The α diversity was calculated considering Shannon and Simpson Indexes and richness was analyzed by Chao1, both calculated in Phyloseq Package on R Environment (Mcmurdie and Holmes 2013). The Permutational Analysis of Variance was performed using function Adonis (Oksanen 2010), with Bray Curtis distance matrix and 999 permutations in order to identify the contribution of the factors (Plant and Time) in the fungi community structure. Principal coordinates analysis (PCoA) was performed to summarize the variation of the community structure in each plant (colours) and time (shapes). Bar plots with the dominant taxonomic groups, (i.e., ten most abundant phyla) were generated to identify the contribution of different Phyla in each treatment (Wickham 2009). The ternary plots were created with Package ggtern (Hamilton and Ferry 2018), considering the relative average of each Phylum.

Core communities are described as the organisms that are widely distributed in a determined environmental type despite variations within it (Hanski 1982) and at the same time, variable communities are organisms which appearance varies along with a given environmental parameter. The identification of the core and variable communities was made by a probabilistic method as described by Gumiere, 2018 (Gumiere et al. 2018) using the R software version 3.2.2 (http://www.r-project.org/). The size of the core and variable communities and also the taxonomic compositions were verified using bar plot graphics. Similarly, the identification of specialists and generalists were performed using the EcolUtils R package (Salazar 2019) according to Levin’s definition of niche breadth (Levins 1968) and 100 permutations. FunGuild (Nguyen et al. 2016b) was used to assign fungal phylotypes to a functional guild where possible.

### 2.6. Nucleotide sequence accession numbers

Fungal ITS rRNA gene sequences obtained in this study are publicly available in the NCBI server under the SRA accession number: SRP151571.

## 3. Results

### 3.1. Alpha and Beta-diversity profiles

The alpha diversity metrics analysis indicated that CHAO1 richness estimator did not differ significantly along time for all the three studied plant species, but in T7 it was lower in *Avicennia shaueriana* compared to the other species (Fig. 1 a). The Shannon diversity index also did not show significant differences along time for *A. shaueriana* and *L. racemosa*; however, *R. mangle* showed a drop on T15 followed by a new increase in diversity in T30 and T60. *L. racemosa* showed highest Shannon diversity compared to the other species mainly on T7 and T30 (Figure 1 b). Simpson index showed the same trend (Figure 1 c).

**Figure 1:**
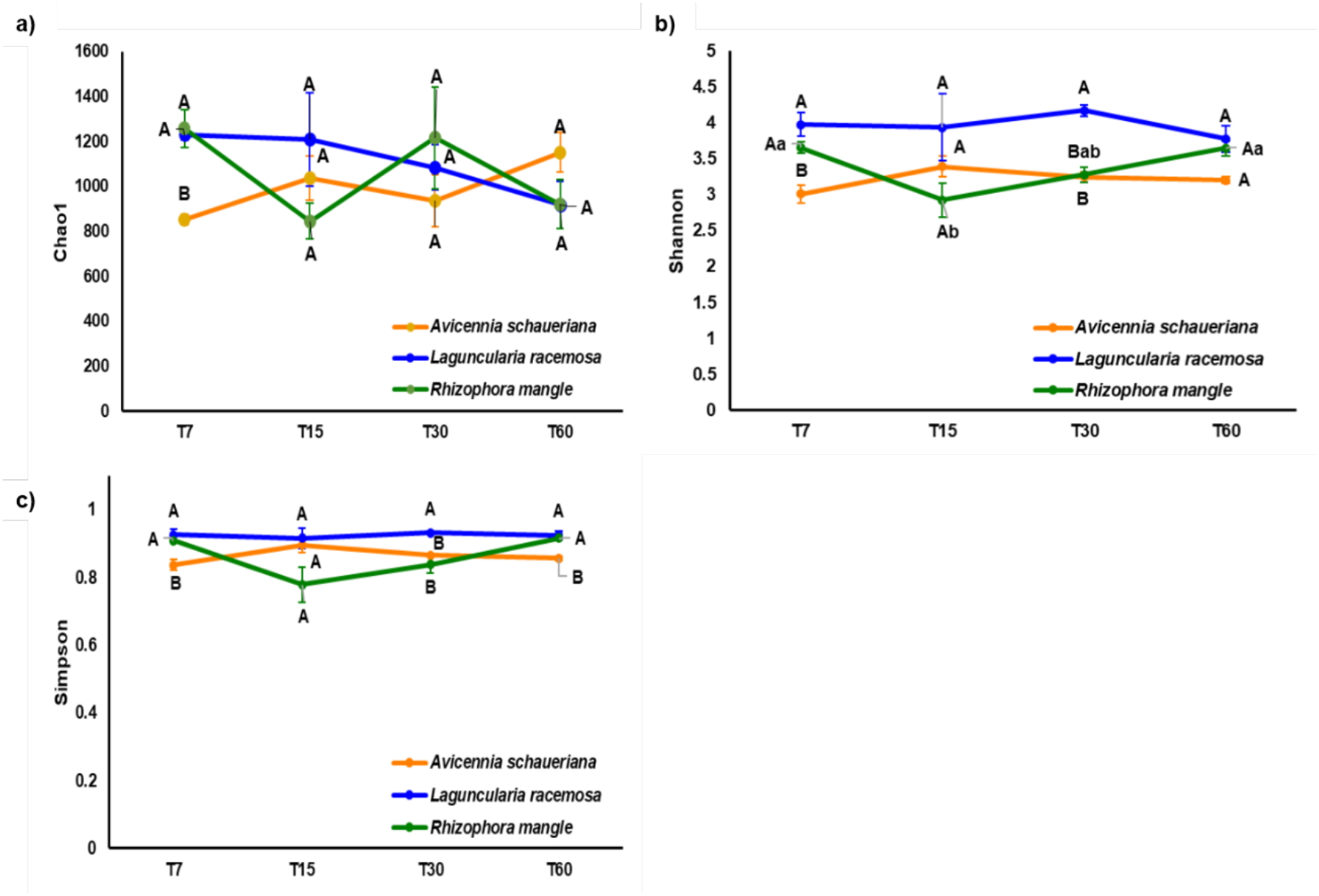
**a)** Chao Index of richness. **b)** Shannon and **c)** Simpson alpha diversity indexes of the fungi community during the decomposition process of three plant species in Cananéia mangrove.

The role of the plant species as the main factor contributing to the fungal community structure during the decomposition process is also evident in the Principal Coordinated analysis (PCoA, ANOVA p<0.05). Clear separation of the samples based on plant material were observed despite no separation of samples based on time (Figure 2 a). The compositional plot of PCoA shows a higher number of OTUs from the phyla Ascomycota and Basidiomycota in all plant species (Figure 2 b). The Permanova analysis showed that plant material was the most significant factor affecting the plant decomposition mediated by the fungi community in Cananéia mangrove (R^2^= 0.41, p=0.001) with time also being slight significant for the process (R^2^= 0.07 considering p<0.1).

**Figure 2:**
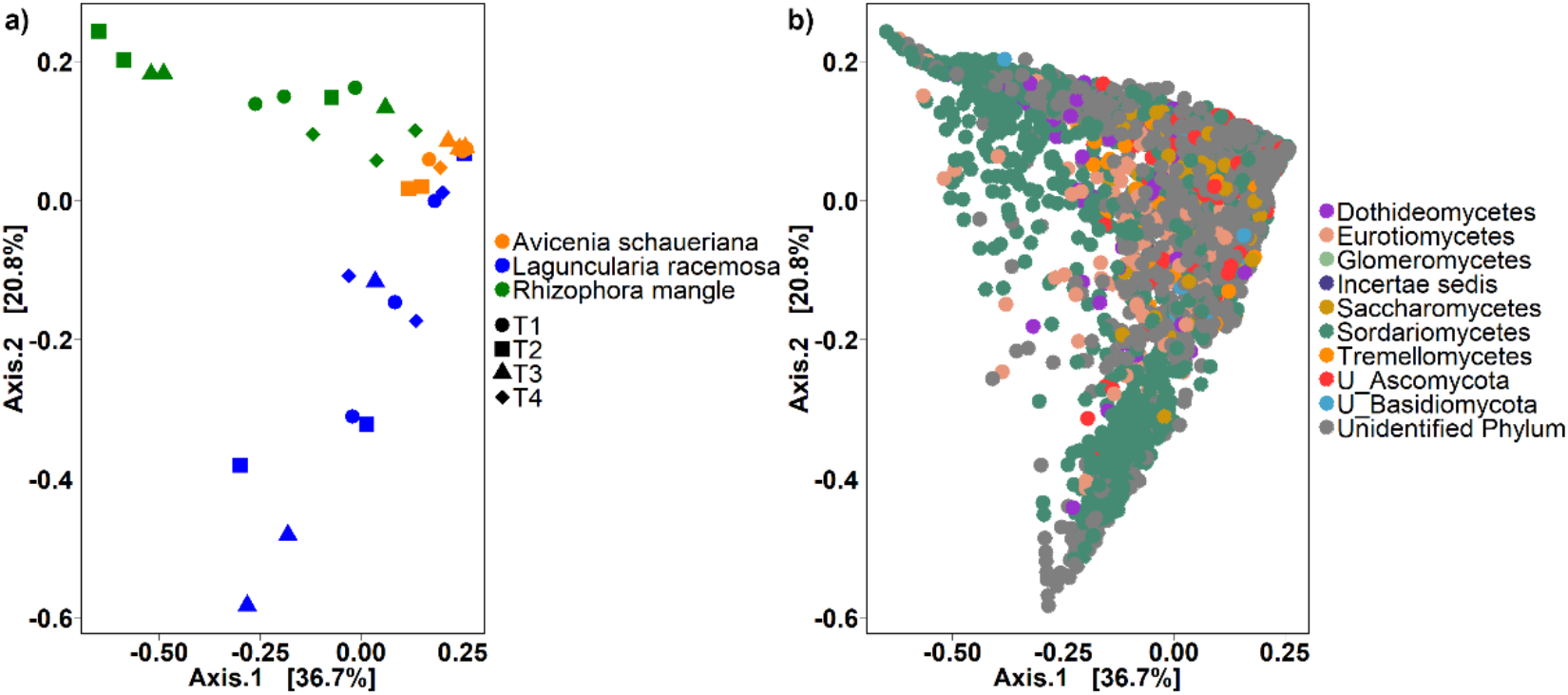
The most abundant fungi families assembled in the three plant species (*Avicennia schaueriana, Laguncularia racemosa and Rhizophora mangle*) during the decomposition process. **a)** Principal Coordinated analysis (PCOa) showing the β diversity of the six most abundant families in the three plant species in the four collection times (7, 15, 30, 60). The first axis explained 38.2% of the data while the second axis explained 23.4%. **b)** Principal Coordinated analysis showing the β diversity of the six most abundant phyla in the three plant species where each dot represent an OTU colored according to the phylum level of the taxonomic classification

### 3.2. Community composition

The six predominant fungal phyla in the three plant species at Cananéia mangrove were *Ascomycota*, *Basidiomycota*, *Chytridiomycota*, Glomeromycota, Incertae sedis and Zygomycota. The plant material from different species selected different taxa during the process of degradation. The most abundant classes common to all the three plant species were Saccharomycetes, Sordariomycetes, Tremellomycetes, Eurotiomycetes, Dothideomycetes, as well as unidentified classes belonging to the Phylum Ascomycota and fungi from unidentified Phylum (Figures 3).

**Figure 3:**
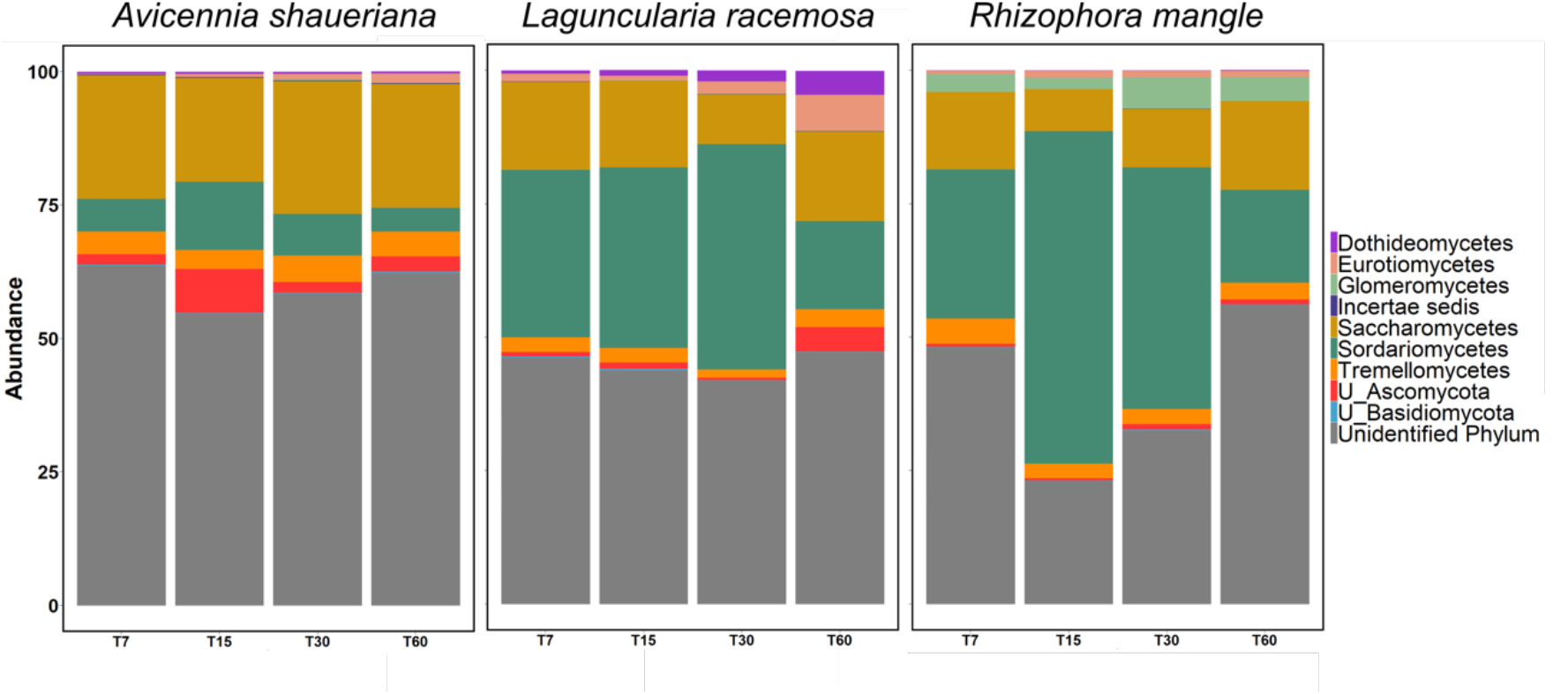
Relative abundance of the 10 most common classes along with the two unclassified groups (U_) belonging to the phyla Ascomycota and Basidiomycota in the *A. schaueriana* plant along the 60 days of decomposition process.

Saccharomycetes was the predominant class in all four stages in *A. shauerianna*. After 15 days, this group presented a drastic reduction while there was a considerable increase in Sordaryomicetes and the unclassified groups of the Phylum Ascomycota. In the samples that followed, Eurotiomycetes and other minor classes presented an increase in abundance. The decomposition process in *L. racemosa* was more dynamic in the final phases (time 30 and 60). Sordariomycetes was the most abundant class until the 30th day. The initial stages (7 and 15 days) presented similar abundances of the classes Sordariomycetes, Saccharomycetes, Tremellomycetes and unclassified classes from the Phylum Ascomycota. In the samples from the 30th day, Sordariomycetes and Eurotiomycetes increased their abundance while Saccharomycetes and Tremellomycetes decreased. In the last sampling time (60th day), the most abundant classes (Sordariomycetes and Saccharomycetes) had very similar proportions while Unclassified Ascomycota, Eurotiomycetes, Dothideomycetes and Incertae sedis had an increase in their proportions compared to the former samples. Similarly, Sordariomycetes was the most abundant in all samples except in the final phase (60th day) in *R. mangle* (Figure 3). This plant also presented a substantial increase in the abundances of Eurotiomycetes during at time 60. Sordariomycetes considerable increased its abundance in time 15 compared to the others. In the Last period, Saccharomycetes and Sordaryomicetes had equal abundances, while Dothideomycetes and unclassified Ascomycota had a slightly increased.

The ternary plots show the distribution of the main OTUs classified at the class level in the three plant species during the four studied phases of the decomposition. The initial stages (Times 7 and 15) (Figure S1 a and b) presented more OTUs corresponding to unclassified classes of phyla Ascomycota and Basidiomycota distributed in the centre with more proximity to the plants *A. schaueriana* and *L. racemosa*. In the intermediary stage (Time 30) (Figure S1 c) most of the OTUs corresponding to the Saccharomycetes e Sordariomycetes were grouped near one side of the triangle corresponding to *A. schaueriana*, with few spaced ones belonging to the classes of U_Basidiomycota near *L. racemosa* and *R. mangle.* The final stage (Time 60) (Figure S1 d) was the one that presented the most homogeneous distribution of the OTUs among all corners of the triangle.

### 3.3. Role of fungi on the community

Unclassified fungi accounted for more than 50% of the organisms in the core community in all the three plant species. The most abundant class in the core community was Saccharomycetes (with more than 10% in all samples) and Sordariomycetes both belonging to the Phylum Ascomycota. Sordariomycetes were expressive more abundant in *R. mangle* (34%) than in *L. racemosa* (8%) and *A. schaueriana* (2%). The phylum Basidiomycota occurred in lower abundance being represented by the class Tremellomycetes (less than 5% for all samples) (Figure 4).

**Figure 4:**
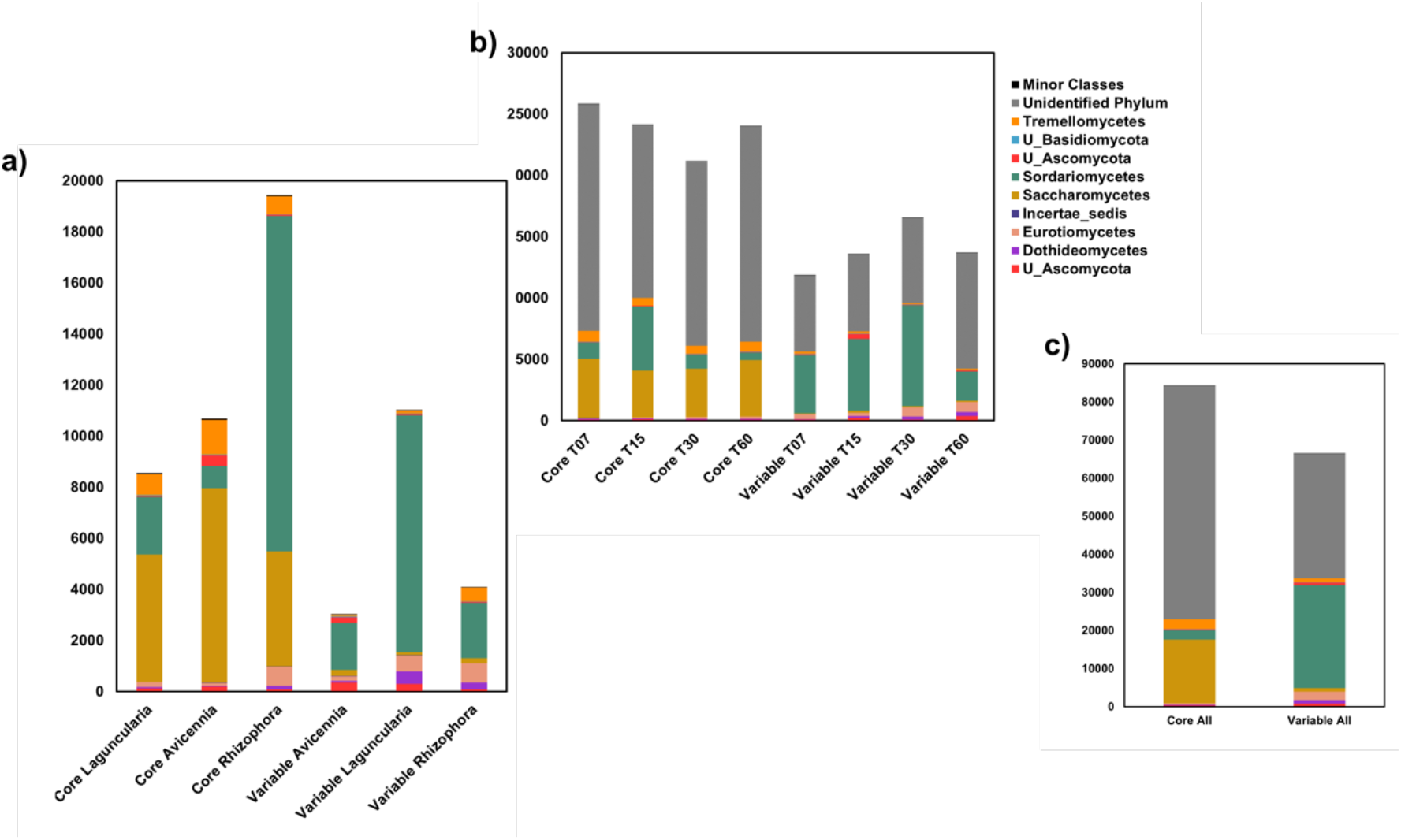
Relative abundance of the OTUs classified as belonging to the core or variable communities determine by the Poisson distribution in the three studies plant species (*Avicennia schaueriana, Laguncularia racemosa and Rhizophora mangle*).

Higher proportions of unclassified microorganisms (more than 50%) also were observed in the variable fungi community followed by the class Sordariomycetes that had higher proportions in *L. racemosa* (40%) than in *A. schaueriana* (16%) and *R. mangle* (17%). As opposed to the core community, the class Saccharomycetes occurred in lower abundance in the variable communities (less than 3% for all samples). The other variable classes that composed the variable communities were Eurotiomycetes, Dothideomycetes (phylum Ascomycota) and Tremellomycetes (phylum Basidiomycota) (Figure 4).

Despite the variation in the frequencies of core (or variable) OTUs over time, some classes were more abundant in one group rather than the other. Saccharomycetes, Tremellomycetes, and Sordariomycetes were dominants in the core group, while the Sordariomycetes, Eurotiomycetes, Dothideomycetes and unclassified classes were abundant in the variable group.

The separation of OTUs into generalists and specialists (*sensu* Levins), indicates that there were more OTUs that could be significantly (p<0.05) assigned to the latter than the former (figure 5). Saccharomycetes were the most abundant in both groups, followed by Sordariomycetes. However, within the specialists, the Sordariomycetes was less abundant in *A. shaueriana* than the other plant material. Also, Tremellomycetes was the third most frequent class among the generalists and was almost undetected among specialists indicating that most of the individuals of this class found in this samples must have a large niche breadth.

**Figure 5:**
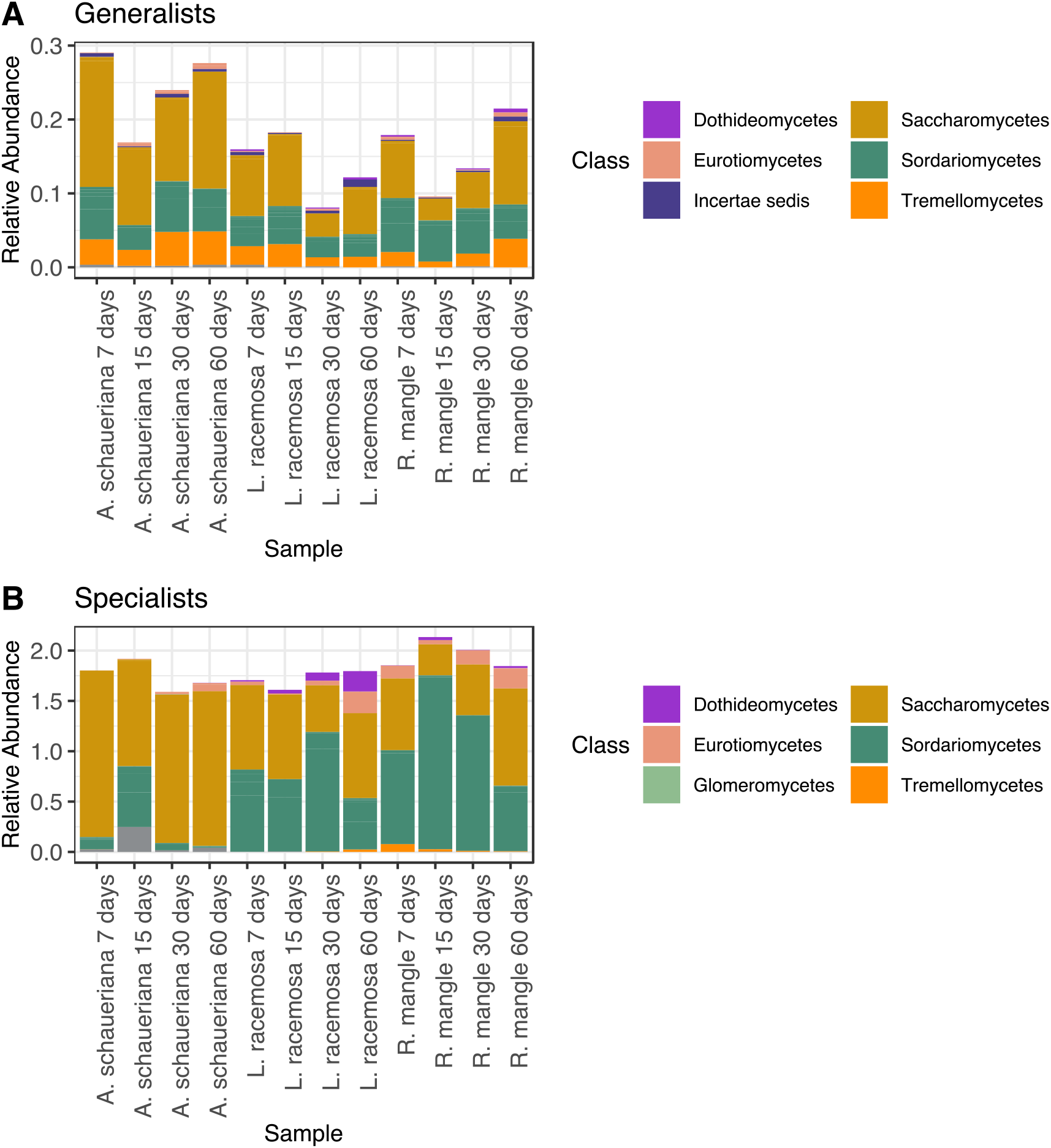
Taxonomic classification of the fungal OTUs identified as Generalists or Specialists found during the decomposition of mangrove leaves.

In terms of relative abundance, *A. shaueriana* decaying leaves had the highest abundance of generalists than the other material while the specialist’s frequency was more stable. Also, a temporal variation could be observed for the former while the latter presented low variation. The three plant materials showed a high abundance of generalists the seventh day (and fiftieth day for *L. racemosa*), with a sharp decline after that and a steady increase in the last two samples.

The identification of the functional guilds to which the detected OTUs were assigned (Nguyen et al. 2016a) also indicated that a difference among the different decaying material and time of sampling (figure 6). *A. shaueriana* had an overall higher proportion of saprotrophs from different groups than the remaining leaves. Interestingly, the other plant material had a high proportion of OTUs assigned as a plant pathogen. Most of these OTUs were identified as belonging to the genus *Pilidiella* and *Pestalotiopsis*. The *Pilidiella* was only identified in *L. racemosa* while *Pestalotiopsis* was detected in both.

**Figure 6:**
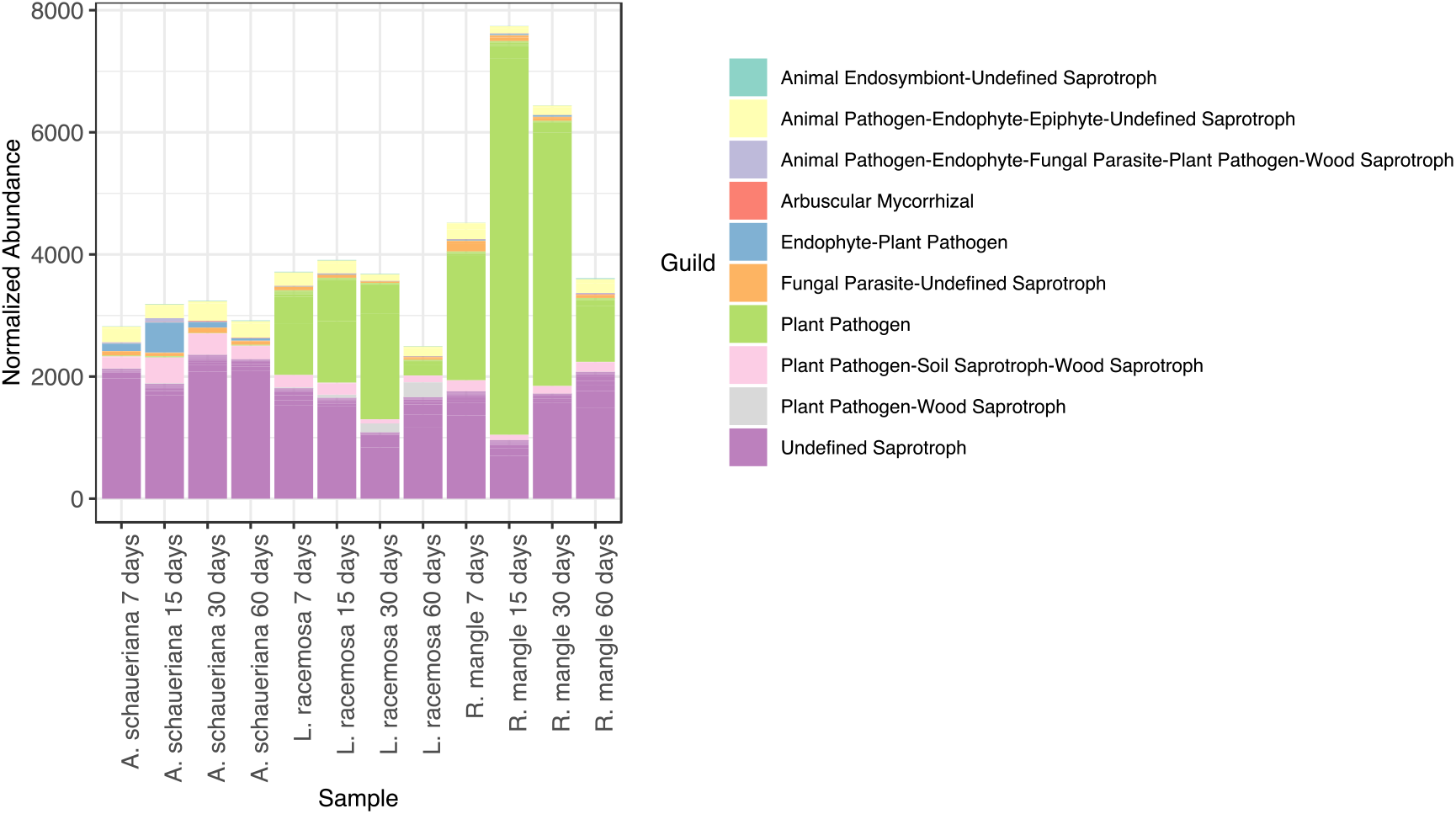
Identification of **OTUs belonging to fungal** guilds found during the decomposition of mangrove leaves. Identification was performed using FunGuild.

## 4. Discussion

In this study, we aimed to provide a phylogenetic analysis of the fungal community involved in leaves decomposition from three mangrove plant species, in Cananéia mangrove, Brazil, for two months. The molecular identification revealed that the decomposition process in the Cananéia mangrove is composed of a diverse community of fungi, and this result is following previous studies of the fungal diversity in this environment (Sebastianes et al. 2013).

Previous studies demonstrated the succession process of fungal communities during decomposition of leaf litter material from different plants and different locations like parks vegetation (Kodsueb et al. 2008; Promputtha et al. 2008); woodland stream (Marano et al. 2011); temperate forests (Osono 2002; Voriskova and Baldrian 2013) and tropical seasonal forest (Osono et al. 2009). During the decomposition of plant material in a temperate forest, fungal biomass increased rapidly from low values to a maximum point at the 60th day of the experiment (Voriskova and Baldrian 2013).

The main factor contributing to the fungal community structure during the decomposition process was the plant species. A previous study with the endophytic fungi community from the three mangrove plants characteristic from Cananéia mangrove, plant species was a factor that affected the fungal community diversity (Sebastianes et al. 2013). However, differently from what was observed to the endophyte’s community (Sebastianes et al., 2013), in the decomposition process, *L. racemosa* was the plant that presented the lowest numbers of populations of endophytic fungi, followed by *Rhizophora mangle*. *A. schaueriana* was the plant species with the highest community of endophytic fungi.

Previously studies have already shown that in mangroves, the fungal colonization of plants, the decomposition process, and the succession stages vary according to the kind of mangrove substrate like plant species (Ananda and Sridhar 2004; Costa et al. 2012). Possibly this differences in community composition and assembly are affected by leaf chemical characteristics (Costa et al. 2012). For instance, species of Rhizophoraceae have a high concentration of tannin, a phenolic substance that can inhibit fungal growth (Costa et al. 2012), *A. schaeuriana* present high concentration of salt on their leaves (Sobrado 2004; Dias et al. 2012), while *L. racemosa* presents less salt secretion on their leaves than *A. schaueriana* (Sobrado 2004; Moitinho et al. 2019) and less recalcitrant compounds than *R. mangle* (Barroso-Matos et al. 2012; Moitinho et al. 2018). The chemical content of these plants differs, and this influences how palatable the material is to each microorganism (Barroso-Matos et al. 2012). It also influences the bacterial community composition and succession process (Moitinho et al. 2018).

Core communities are described as the organisms that are largely distributed in a determined environmental type despite variations within it (Hanski 1982); these organisms can be crucial in the maintenance of this habitat. Among the fungal community, the core group of species exerts a significant influence on the turnover of litter in mangrove ecosystems (Ananda and Sridhar 2004). Classes belonging to the Ascomycota were predominant in the core and variable community in the three studied plants, indicating that they can exploit multiple litter types. Saccharomycetes were the predominant class in the core group, with higher concentrations in the *R. mangle* plant compared with the two other plants, while in the variable community Sordariomycetes were more abundant, mainly in the *L. racemosa* plant. Indicating that these groups have some degree of specificity to the plant substrate.

There is a conceptual difference between core taxa and specialist (as well as a variable and a generalist). While the core is an OTU that is expected to be found in every sample of a given treatment (as opposed to the variable that is not, (Gumiere et al. 2018)), a specialist is an OTU that has a smaller niche breadth (NB) that a determinist (Levins 1968). Thus, the characteristics of the environment, such as the resources present, would lead be key to determining ones NB. Therefore, the detection of more generalist taxa in *A. shauerianna* leaves indicates that either there is a higher abundance of different resources or that they are more labile. This observation is supported by the faster degradation rates of *A. shauerianna* (Sessegolo and Lana 1991). This might also be the reason for the higher abundance of Sordariomycetes in the specialists’ group *L. racemosa* and *R. mangle*. These OTUs might be related to the adapted or degrade recalcitrant materials such as lignin and tannins that are present in higher concentrations in these plants (Sessegolo and Lana 1991).

The Phylum Ascomycota is an essential group in the initial stages of the decomposition process of plant material in different environments (Voriskova and Baldrian 2013; Miura et al. 2015; Behnke-Borowczyk and Wołowska 2018). They are one of the dominant phyla of fungi in Brazilian mangroves (Sebastianes et al. 2013) and were predominant in the rhizosphere and bulk soil collected from the Red Sea grey mangroves (*Avicennia marina*) (Simões et al. 2015). Members of this Phylum generally decompose cellulose selectively over lignin (Osono 2007) and dominate the fungal community and are highly active (production of extracellular enzymes) in the early stages of litter decomposition (Zhang et al. 2018). Aquatic ascomycetes play important roles in wood and leaf litter degradation since basidiomycetes are found in lower quantities (Valderrama et al. 2016). Ascomycetes appear to be better able to withstand the conditions prevailing in aquatic habitats, far more than basidiomycetes (Jones and Choeyklin 2007).

The phylum Basidiomycota was described in mangrove ecosystems in lower quantities than the Ascomycota phylum, which corroborates with the results found in this study (Sebastianes et al. 2013). The phylum Basidiomycota also is constant during the decomposition process of leaf material, and are considered to have more ligninolytic abilities than the ascomycetes (Osono 2007); and are correlated with the late stages of decomposition of leaf litter (Osono 2007; Voriskova and Baldrian 2013).

The main classes of fungi found in this study were previously reported in different decomposition studies from different environments. Leotiomycetes, Sordariomycetes and Eurotiomycetes were the abundant classes during decomposition of litter from a forest of Austria (Schneider et al. 2012). Sordariomycetes, Saccharomycetes, Eurotiomycetes and Dothideomycetes classes were described during the decomposition process of wheat straw (Zhang et al. 2014). Ascomycota, Dothideomycetes, and Leotiomycetes dominated in the communities, and there was a high relative abundance of Sordariomycetes. They were also the most abundant groups on submerged litter decomposition in streams (Koivusaari et al. 2019). Sordariomycetes also was the most recovered class from aquatic environments from Mexico and the other classes like Saccharomycetes, Eurotiomycetes. Leotiomycetes, Dothideomycetes, Sordariomycetes, Tremellomycetes and Chytridiomycetes also were observed in this study (Valderrama et al. 2016). Microbotryomycetes are Basidiomycete yeasts common to aquatic environments like mangroves. Agaricomycetes also were isolated from mangrove wood substrate and Incertae sedis are common fungi in decaying leaf litter in streams (Jones and Choeyklin 2007). Chytridiomycota also is an abundant phylum described in aquatic environments (Valderrama et al. 2016).

The high abundance of saprotrophic OTUs in this material confirms that the taxa found are most likely involved in the degradation of the leaf material. It also indicates a lower abundance of cross feeders. However, the high abundance of OTUs related to the plant pathogenic *Pilidiella* and *Pestalotiopsis* fungi indicates that some abundant OTUs might not be active in the degradation of these materials. Nevertheless, despite the presence of many plant pathogens in these genera, several isolates were shown to degrade lignin (Arfi et al. 2013; Raudabaugh et al. 2018).

Altogether, our results indicated that the different plant materials selected for specific fungal communities. However, there was a dominance of Ascomycota and later an increase in the abundance of Basidiomycota. This change reflected in the differences of the core and variable community between plant materials and overtime, where we observed a similar contrast in the weight of each group in the entire community.

In conclusion, the results presented in this work show that there is a specialization of the fungal community while degrading different plant species of mangrove. This is associated with the different rates of degradation expected by the chemical composition of the leaves. For instance, OTUs such identified as Saccharomycetes are more abundant and have in samples of *A. shauerianna* with lower concentrations of tannins and lignin. It also provides essential information about the frequency and composition of the main fungal classes of the communities associated with the degradation of three plants in the Brazilian mangrove forests.

## Author contributions

**Marta A. Moitinho:** Conceptualization; Data curation; Formal analysis; Investigation; Methodology; Software; Roles/Writing - original draft; Writing - review & editing; **Josiane B. Chiaramonte;** Data curation; Formal analysis; Investigation; Methodology; Software; Visualization; Roles/Writing - original draft; Writing - review & editing; **Laura Bononi;** Methodology; Roles/Writing - original draft; Writing - review & editing; **Thiago Gumiere:** Conceptualization; Data curation; Formal analysis; Methodology; Software; Supervision; Visualization; Roles/Writing - original draft; Writing - review & editing; **Itamar S. Melo:** Conceptualization; Project administration; Resources; Supervision; Validation; Visualization; Roles/Writing - original draft; Writing - review & editing; **Rodrigo G. Taketani:** Conceptualization; Data curation; Formal analysis; Funding acquisition; Investigation; Project administration; Resources; Software; Supervision; Validation; Visualization; Roles/Writing - original draft; Writing - review & editing;

