## Supplementary material for "Fungal succession on the decomposition of three plant species from a Brazilian mangrove": figure S1

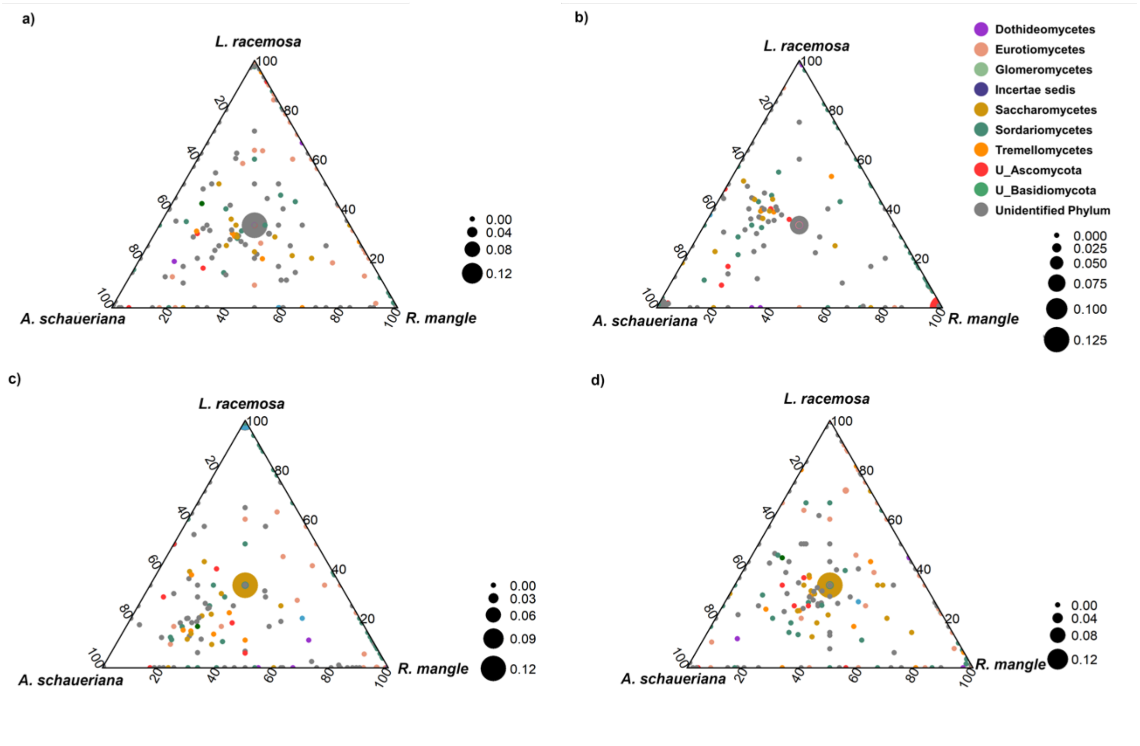


**Figure S1**: Ternary plots of the distribution of the OTUs across the three plant species (*A. schaueriana, L. racemosa* and *R. mangle*) along different times of the decomposition (**a)** time 7. **b)** time 15, **c)** time 30 and **d)** time 60). Each point represents an OTU, and its position indicates the proportion of its relative abundance at the different plant species. Points closer to the ternary plot corners indicate that a greater proportion of the total relative abundance of this OTU was found in this particular environment. Point colors indicate the phylum of the OTU. The lines inside the ternary plot indicate the X level of relative abundance of each of the sites. Only the OTUs with a per-site mean relative abundance of more than 0.1% are shown.
